# C1q-mediated steric hindrance limits FcγR contribution to antibody-driven phagocytosis by human neutrophils

**DOI:** 10.64898/2026.09.30.755584

**Authors:** Max L.B. Grönloh, Lisanne de Vor, Seline Zwarthoff, Jana Venema, Rosemieke Smits, Stan Baijens, Maartje Ruyken, Piet Aerts, Carla J.C. de Haas, Kok van Kessel, Leendert A. Trouw, Bart Bardoel, Suzan Rooijakkers

**Author notes:** These authors contributed equally (Max L.B. Grönloh, Lisanne de Vor). Corresponding authors: Suzan Rooijakkers & Max L.B. Grönloh.

## Abstract

Phagocytosis and killing of bacteria by neutrophils is a key driver of human immunity against invading infections. IgG antibodies targeting the bacterial surface can potently enhance phagocytic uptake by engaging Fc gamma receptors (FcγRs) on the neutrophil membrane. In addition, IgG-mediated complement activation triggers surface deposition of complement opsonins that engage complement receptors (CRs) to further enhance target cell clearance. In this paper, we reveal that complement deposition prevents engagement of FcγRs to the Fc-domain of surface-bound IgG molecules. When complement is present, we observe that blockade of FcγRs does not influence phagocytosis of both Gram-positive and -negative bacteria (using *Staphylococcus aureus* and *Klebsiella* pneumoniae as model organisms). Complement-mediated hindrance of FcγR engagement was observed in different model conditions, for example using either pre-existing polyclonal antibodies in human serum or recombinant monoclonal antibodies (mAbs) targeting different bacterial antigens. Using Fc engineering, we found that inhibition of FcγR binding is dependent on the ability of IgGs to form IgG hexamers. Mechanistically, we show that complement proteins, in particular C1q, directly prevents the interaction between IgG-Fc and FcγR by steric hindrance. Together, these insights help us define the respective roles of FcγR and CR in neutrophil phagocytosis, while also providing important fundamental insights for therapeutic antibody development.

**Significance Statement:** Antibodies can recruit immune cells through Fc receptors or activate complement, but how these pathways interact when both are engaged on the same target remains poorly understood. We show that complement, and specifically C1q, can physically restrict access of Fcγ receptors to antibody Fc domains, shifting immune-cell recognition toward complement receptors. This mechanism limits the contribution of Fcγ receptors to neutrophil phagocytosis of bacteria and has important implications for antibody engineering, where complement and Fc-receptor functions are often optimized simultaneously. More broadly, our findings reveal a previously underappreciated mechanism of crosstalk between two major arms of antibody-mediated immunity, with potential relevance to antibacterial, antiviral, and anticancer antibody responses.

## Introduction

Phagocytosis is the principal line of host defense against extracellular bacteria by professional phagocytes, most notably migrating neutrophils and tissue-resident macrophages. Efficient bacterial clearance requires a coordinated process involving recognition of target cells via membrane receptors, internalization and intracellular destruction. While phagocytosis can be initiated through pattern recognition receptors (PRRs) that recognize conserved microbial structures, opsonization by immunoglobulins (Igs) and complement proteins greatly enhances the phagocytic process. Phagocytes possess membrane-associated receptors against both Igs and complement proteins, namely Fc receptors and complement receptors (CRs).^1^

Among Fc receptors, the most studied is the family of Fc gamma receptors (FcγRs), which recognize IgGs through their constant (Fc) region. FcγRs are expressed by a wide range of immune cells, including neutrophils, monocytes and macrophages. Besides triggering phagocytosis, FcγRs are relevant in antibody-dependent cell-mediated cytotoxicity of cancer cells and the release of inflammatory cytokines.^2^ Human neutrophils express FcγRIIa (CD32) and FcγRIIIb (CD16b) constitutively, and upregulate FcγRI (CD64) under certain inflammatory conditions.^3^

CRs, including CR1 (CD35) and CR3 (CD11b/CD18) expressed by neutrophils, are cell-surface receptors that recognize complement opsonins deposited on microbial surfaces. One route of complement deposition is IgG-mediated activation of the classical complement pathway, which is initiated when C1q binds to noncovalently assembled IgG hexamers.^4^ This interaction activates a proteolytic cascade that first results in the covalent deposition of C4b on the bacterial surface. Surface-bound C4b subsequently contributes to formation of the classical-pathway C3 convertase, which cleaves C3 and promotes the extensive deposition of C3b. C3b can then be converted into its derivative iC3b, which like C3b also is recognized by CRs.^5^ As C3 fragments densely decorate the microbial surface, CRs are a highly efficient mechanism for pathogen recognition and phagocytosis.

The growing burden of antimicrobial resistance,^6^ together with the vulnerability of immunocompromised patients to invasive bacterial infections,^7^ has increased interest in antibody-based therapies that can complement conventional antibiotics. Despite the success of monoclonal antibodies (s) against viral infections,^8,9^ antibody development for bacterial infections has progressed more slowly.^10^ Whereas antiviral antibodies can often provide protection by directly neutralizing viral particles and preventing host-cell entry, antibodies against surface structures intended to promote the elimination of intact extracellular bacteria frequently additionally rely on immune effector mechanisms. Their activity therefore depends not only on antigen specificity and affinity, but also on their ability to engage FcγRs and activate complement, thereby promoting recognition through CRs. To date, the clearest clinical successes of antibacterial mAbs have involved the neutralization of secreted toxins,^10–12^ whereas antibodies designed to promote the clearance of intact bacteria have shown more limited clinical efficacy.^10,13,14^

*Staphylococcus aureus* and *Klebsiella pneumoniae* are clinically important causes of bloodstream infections, pneumonia, and other invasive infections,^6^ and antibiotic-resistant lineages of both pathogens have been prioritized for the development of new therapeutic strategies.^15^ *S. aureus* is a Gram-positive pathogen that expresses numerous surface proteins and cell-wall glycopolymers while employing extensive mechanisms to evade antibody- and complement-mediated immunity.^16–18^ *K. pneumoniae* is an encapsulated Gram-negative opportunistic pathogen whose capsule and lipopolysaccharide contribute to complement resistance and impaired phagocytic recognition.^19,20^ mAbs against toxins, surface proteins, capsular polysaccharides and other cell-surface glycans of these pathogens have been characterized.^21–25^ These antibodies provide valuable tools for determining how antigen specificity and opsonization context control complement activation, phagocytosis, and bacterial killing. However, when antibody Fc domains and complement-derived opsonins are simultaneously present on the bacterial surface, the relative contributions of FcγRs and complement receptors remain poorly understood.

Here, we investigated the relative contributions of FcγRs and complement receptors to the phagocytosis and killing of *S. aureus* and *K. pneumoniae* by human neutrophils. We show that phagocytosis and intracellular killing are largely maintained in the presence of complement following FcγR inhibition, indicating a dominant role for complement receptors under these conditions. Mechanistically, we demonstrate that complement deposition, and specifically C1q, prevents access of FcγRs to IgG-Fc domains on the bacterial surface, thereby functionally limiting FcγR engagement. Together, our findings reveal that complement not only enhances opsonization but also suppresses FcγR-mediated recognition, redefining the interplay between these two central pathways in antibody-dependent immunity.

## Results

### Blocking FcγRs does not impair phagocytic uptake in the presence of complement

First, we aimed to study the relative contributions of FcγRs and CRs in phagocytic uptake and bacterial killing of *S. aureus*. To account for the influence of staphylococcal immune evasion proteins that inhibit IgG-Fc interactions, we used the *S. aureus* LAC strain lacking Protein A (Spa) and Second Immunoglobulin-binding protein (Sbi). (LAC Δ*spa/sbi*)^16,26^ As a source of complement, we used pooled serum from healthy human donors, which also contains substantial levels of naturally acquired antibodies against *S. aureus*.^27^ To not engage CRs, we heat-inactivated the serum. The efficiency of this approach was confirmed by a clear loss of C1q and C3b deposition on *S. aureus* in conditions with heat-inactivated serum (Figure S1A-B). Inhibition of FcγR-IgG interactions was achieved by treating neutrophils with Formyl Peptide Receptor-like 1 inhibitory protein-like (FLL/denoted as FcγR inhibitor).^28,29^ Next, we co-incubated GFP-expressing *S. aureus* together with neutrophils and measured phagocytosis by measuring GFP signal of neutrophils via flow cytometry. Strikingly, no decrease in phagocytosis was detected when inhibiting FcγRs in the presence of complement, while FcγR-mediated phagocytosis could be fully blocked when complement components had been thermally inactivated (Figure 1A-B). When comparing both receptor pathways in isolation, it stands out that CR-mediated phagocytosis of *S. aureus* in human serum leads to significantly more phagocytosis compared to FcγR-mediated phagocytosis in heat-inactivated serum (Figure 1A-B).

**Figure 1.**
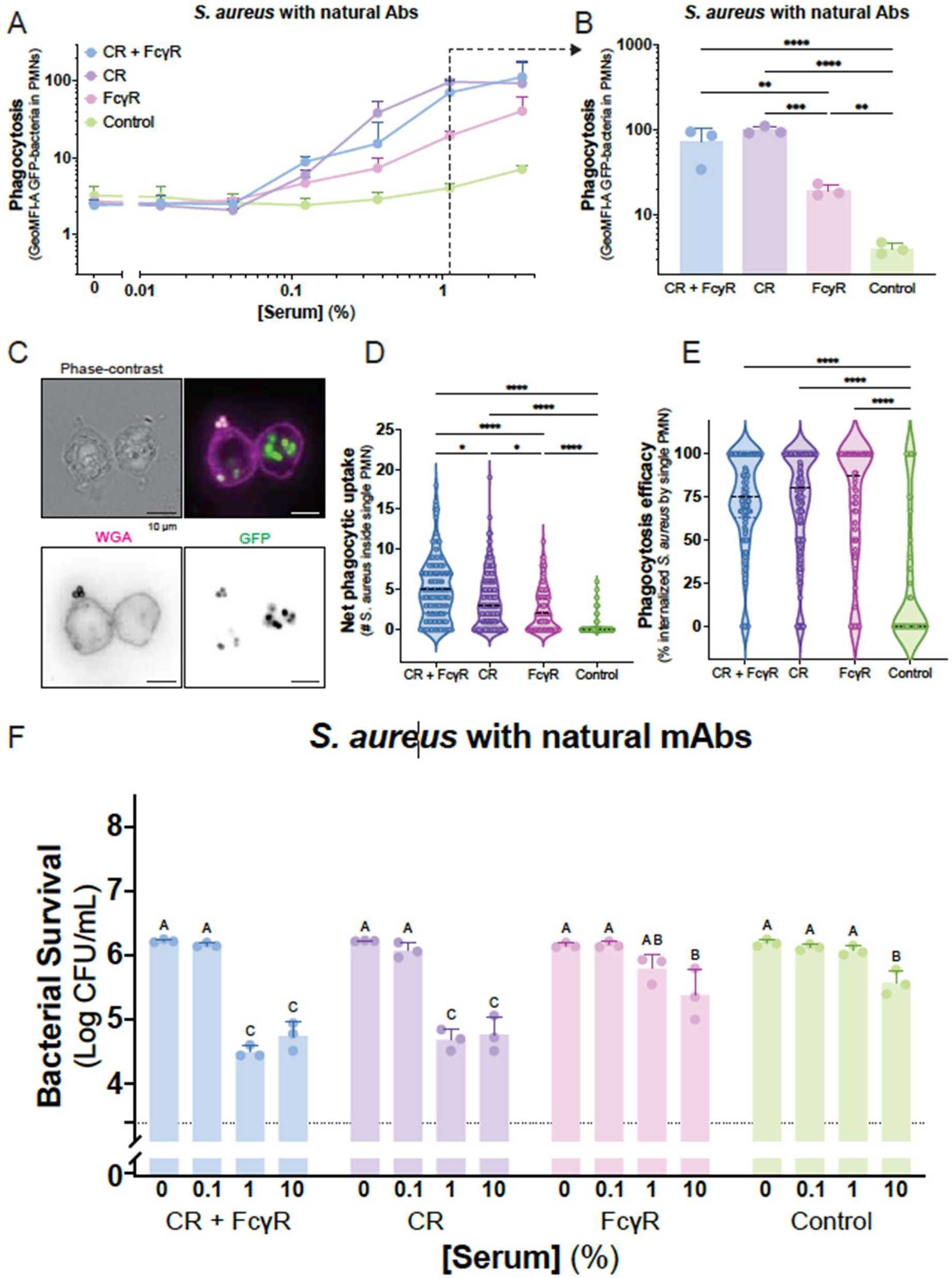
Blocking FcγRs in the presence of complement does not impair phagocytic uptake or bacterial killing. **(A-F)** (Heat-inactivated) human serum-dependent phagocytosis of *S. aureus* LAC Δ*SpA/Sbi* - GFP by human neutrophils (MOI 10:1). Neutrophils were pre-incubated with 3 µg/mL FLIPr-like to inhibit FcγRs or buffer and were allowed to phagocytose opsonized bacteria for 15 minutes at 37 °C, after which **(A-B)** GeoMFI of GFP in the neutrophil population was determined via flow cytometry or **(C-F)** intra- and extracellular bacteria per neutrophil was measured via widefield microscopy. **(A)** (Heat-inactivated) human serum concentration-dependent phagocytosis. Data represent mean ± SD of n=3. **(B)** Phagocytosis at 1.1% (heat-inactivated) human serum. Data represent mean ± SD of n=3. A One-way ANOVA test was used to determine significant difference between the indicated bar graphs in which p values are indicated with ∗∗p < 0.01, ∗∗∗p < 0.001, and ∗∗∗∗p < 0.0001. **(C-D)** Phase-contrast and widefield fluorescent (green: GFP, magenta: WGA) images. Orange triangle indicates extracellular *S. aureus* and blue triangle indicates intracellular *S. aureus.* Scale bar is **(C)** 10 µm / **(D)** 25 µm. **(E-F)** Quantification of **(E)** net phagocytic uptake (number of internalized *S. aureus* and **(F)** phagocytic efficacy (number of internalized *S. aureus* divided by sum of internalized and extracellular *S. aureus*). Dots represent individual neutrophils, data are represented by violin plots in which the medians are displayed by a dotted line. **(E-F)** A Kruskal-Wallis test was used to determine significant difference between the conditions in which p values are indicated with p < 0.05, ∗∗∗p < 0.001, and ∗∗∗∗p < 0.0001. **(G)** Heat-inactivated) human serum bacterial viability of *S. aureus* LAC Δ*SpA/Sbi* - GFP after co-incubation with human neutrophils (MOI 1:1). Neutrophils were pre-incubated with 3 µg/mL FLIPr-like to inhibit FcγRs or buffer and were allowed to phagocytose opsonized bacteria for 90 minutes 37 °C at 5% CO_2_. Data are visualized as log-transformed CFU/mL values, and displayed as mean ± SD of n=3. Statistical significance was assessed by two-way ANOVA followed by Tukey’s multiple-comparisons test. Groups that do not share a letter are significantly different (p < 0.05). **(A-F)** Condition legend: ‘CR + FcγR’ = human serum + neutrophils, ‘CR’ = human serum + FLIPr-like incubated neutrophils, ‘FcγR’ = heat-inactivated human serum + neutrophils, ‘Control’ = heat-inactivated human serum + FLIPr-like incubated neutrophils.

Since our flow cytometry setup does not allow the discrimination between intra- and extracellular bacteria, we developed a microscopy-based approach to verify that FcγR-inhibition in the presence of complement does not impact phagocytosis. Briefly, wheat germ agglutinin (WGA) was used to stain neutrophil membranes as well as extracellular *S. aureus* bacteria. Since WGA does not cross the neutrophil plasma membrane, intracellular engulfed *S. aureus* stays WGA-negative (Figure 1C). Using this approach, we observed that neutrophils can easily phagocytose multiple bacteria, with occasional cells containing over 15 bacteria (Figure 1D & S2A-B). Consistent with the flow cytometry data we show that CR-mediated phagocytosis results in higher net uptake compared to FcγR-mediated phagocytosis (Figure 1). Interestingly, FcγR inhibition in the presence of complement led to a modest but significant decrease in net uptake, while phagocytic efficacy (fraction of associated bacteria internalized) stayed unaltered (Figure 1D-E). Therefore, differences in net uptake are likely driven by differences in initial bacterial association with neutrophils. Following phagocytic uptake, neutrophils internalize bacteria into phagosomes that fuse with granules containing antibacterial products, resulting in enzymatic degradation of the bacteria, acidification, and increased reactive oxygen species (ROS) production.^3^ We hypothesized that even though FcγR-mediated phagocytosis minimally contributed on the level of phagocytic uptake in the presence of complement, perhaps FcγRs still could work synergistically with CRs on the level of bacterial killing, especially since both receptor types induce different intracellular pathways upon activation.^1^ However, intracellular killing of *S. aureus* largely correlated with the degree of phagocytic uptake. CR-mediated phagocytosis led to much more efficient killing compared to FcγR-mediated phagocytosis, and inhibition of FcγRs did not enhance killing in the presence of complement (Figure 1G). Together, these results showed that inhibition of FcγR-IgG interaction using FLIPr-like reduced phagocytosis potently, but only in the absence of complement proteins. This raises an interesting paradox, in which complement might on one hand boost phagocytosis via but on the other hand potentially diminishes the role of FcγRs.

### Complement-mediated phagocytosis remains dominant across antibody targets and bacterial species

Next, we asked whether the observed effect of polyclonal antibodies is generalizable across mAbs targeting different *S. aureus* and *K. pneumoniae* antigens (Figure 2A). For all mAbs, we selected strains to which we observed potent mAb binding and C3b deposition, and CR- as well as FcγR-mediated phagocytic uptake (Figure S3A-H). Additionally, the *K. pneumoniae* strains we tested are resistant to MAC-mediated killing (Figure S3I-K), allowing us to look at phagocytosis-mediated killing only (Figure S3. For *S. aureus*, we produced three IgG1 mAbs against distinct targets, all isolated from B cells derived from patients recovering from *S. aureus* infections. First, 4461 is an antibody recognizing α-N-Acetylglucosamine (GlcNAc)-modified wall techoid acid (WTA).^22,30^ Similarly, 4497 is specific to β-GlcNAc-modified WTA,^22,31^ while rF1 recognizes O-linked GlcNAc modifications on staphylococcal serine-aspartate repeat (SDR) proteins.^32^ Again, we observed an almost complete reduction of phagocytic uptake in the absence of complement components (Figure 2B-D). Conversely, when performing phagocytosis experiments using antibody-depleted human serum, we observed that inhibiting FcγRs did not reduce phagocytosis in the presence of complement proteins, only measuring a small decrease in the 4461 clone (Figure 2B-D). Overall, bacterial survival of *S. aureus* matched to phagocytic uptake for all three clones (Figure 2E-G).

**Figure 2.**
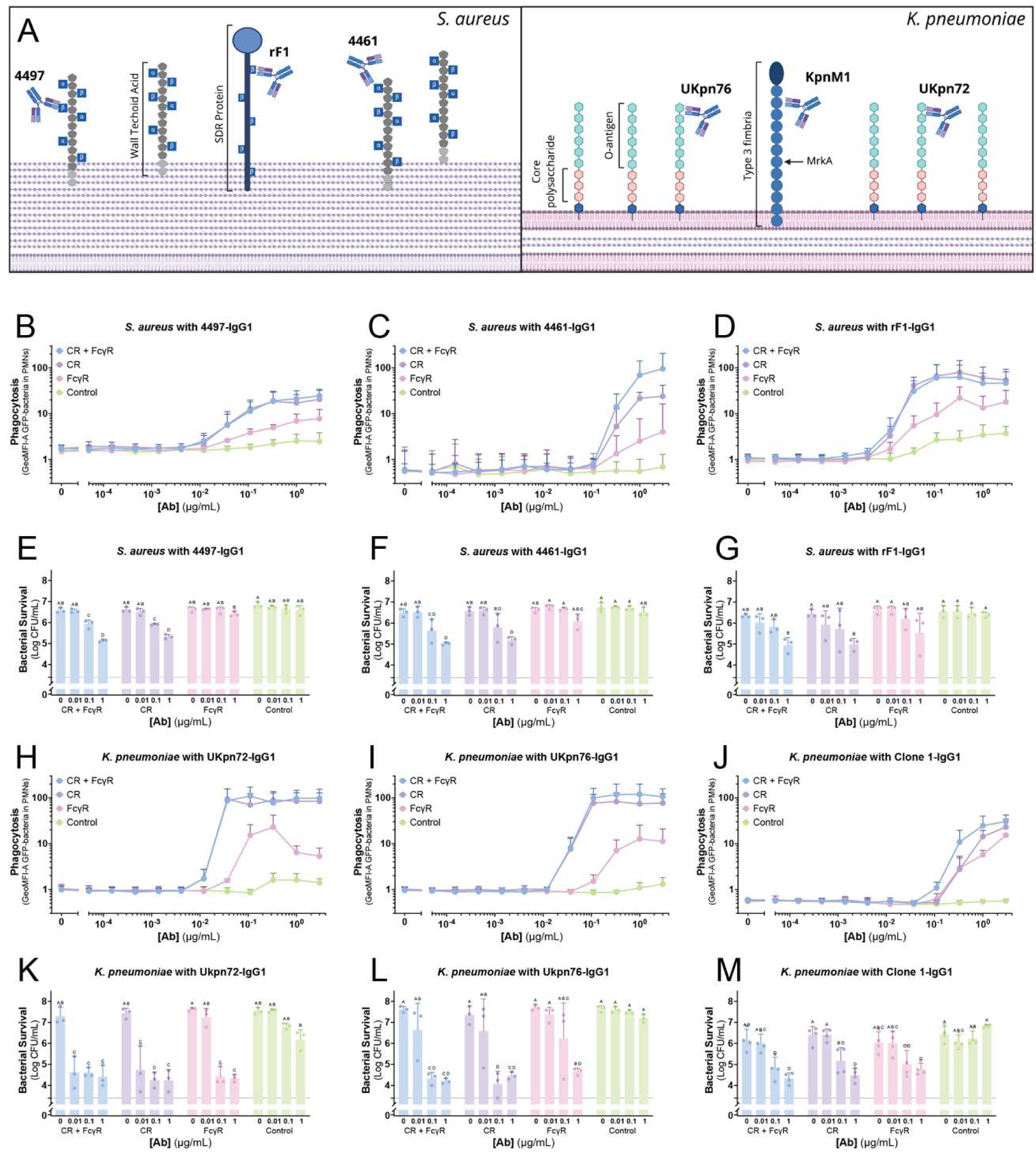
Complement-mediated phagocytosis remains dominant across antibody targets and bacterial species. **(A)** Schematic overview of the mAbs used in this study, together with their targets on *S. aureus* and *K. pneumoniae.* Created in Biorender. **(B-D & H-J)** Phagocytosis (MOI 10:1, 15 min incubation) and **(E-G & K-M)** bacterial survival (MOI 1:1, 90 min incubation) of **(B-G)** *S. aureus* LAC Δ*SpA/Sbi* - GFP, **(H-I & K-L)** *K. pneumoniae Kpn*O1 - GFP and **(J & L)** *K. pneumoniae* Kpn*570.879 -* GFP after coincubation with human neutrophils, which were pre-incubated with 3 µg/mL FLIPr-like to inhibit FcγRs or buffer. Bacteria were opsonized with 1% (*S. aureus)* or 5% (*K.* pneumoniae) IgG/IgM-depleted human serum or buffer, together with a titration of mAb IgG1: **(B & E)** anti-WTA 4497, **(C & E)** anti-WTA 4461, **(D & G)** anti-SDR rF1, **(H & K)** anti-O1-Ukpn72, **(I & L)** anti-O1-Ukpn76, or **(J & M)** anti-MrkA-KpnM1. **(B-D & H-J)** GeoMFI of GFP in the neutrophil population was determined via flow cytometry and data represent mean ± SD of n=3. **(E-G** & K-M) Data are visualized as log-transformed CFU/mL values and displayed as mean ± SD of n=3 (except KpnM1, n=4). Statistical significance was assessed by two-way ANOVA followed by Tukey’s multiple-comparisons test. Groups that do not share a letter are significantly different (p < 0.05). **(B-M)** Condition legend: ‘CR + FcγR’ = IgG/IgM-depleted human serum + neutrophils, ‘CR’ = IgG/IgM-depleted human serum + FLIPr-like incubated neutrophils, ‘FcγR’ = buffer + neutrophils, ‘Control’ = buffer + FLIPr-like incubated neutrophils.

For *K. pneumoniae*, three more IgG1 mAbs were produced. Briefly, we produced KpM1 (*Klebsiella pneumoniae* MrkA antibody clone 1), which targets MrkA, the major component of type III fimbria complex and was generated through a phage screening.^23^ Additionally, we tested UKpn72 and UKpn76, which were derived from B cells from healthy donors and both target the D-galactan II region of the O1-antigen.^24^ For both O-antigen mAbs, FcγR inhibition clearly did not reduce phagocytosis in the presence of complement (Figure 2H-I). This effect was less clearly observed for the MrkA mAb, where the difference between the phagocytosis receptors was not as significant (Figure 2J). For all three clones, treating neutrophils with the FcγR inhibitor did not alter bacterial survival of *K. pneumoniae* (Figure 2K-M). Strikingly different to *S. aureus*, FcγR-mediated phagocytosis of *K. pneumoniae* by itself induced relatively strong intracellular killing in the absence of complement when compared to how many bacteria were taken up by phagocytosis (Figure 2K-M). Overall, we here have demonstrated that, irrespective of antibody specificity, phagocytic uptake of these Gram-positive and Gram-negative species is largely maintained in the presence of complement upon inhibition of FcγRs, suggesting these receptors contribute minimally on the level of phagocytic uptake compared to CRs.

### Inhibition of FcγR-mediated phagocytosis in the presence of complement is independent of IgG subclass

Since IgG subclasses vary in their ability to recruit C1q and activate the complement system,^33–35^ and have different affinities for FcγRs,^36,37^ we subclass-swapped mAb 4497-IgG1 in order to create 4497-IgG2, 4497-IgG3 and 4497-IgG4 variants. All mAbs bound *S. aureus* equally well, while all clones except IgG4 were able to potently activate the complement system, which is in accordance to earlier data from our group (Figure S4A-B).^38^ Irrespective of their subclass, inhibiting FcγRs in the presence of complement proteins had no effect on phagocytosis (Figure S4C-F). As shown in earlier research,^36,37^ IgG2 barely was able to induce phagocytosis in the absence of complement and the small effect that was induced could be completely abolished when adding FcγR inhibitor Figure S4C-F). Compared to IgG1, IgG3 is a more potent in the absence of complement (Figure S4C-F). Especially at high mAb concentrations, FcγR-mediated phagocytosis could not be fully blocked by FcγR inhibitor, preventing us from fully excluding a contribution of FcγR-mediated phagocytosis in IgG3.

### Binding of FcγRIIa to *S. aureus* is reduced by complement deposition

Our results so far led us to hypothesize that complement opsonization could lead to steric hindrance between FcγRs and IgG-Fc. To study FcγR binding to IgG1-Fc in the context of a cell membrane but without disturbance of other opsonophagocytic receptors, we tested binding of antibody-coated bacteria to cell lines overexpressing CR1, CR3, or FcγRIIa, the most prominent FcγR on neutrophils in the context of phagocytosis.^39,40^ CR1- and CR3-expressing cells started binding bacteria around 0.1% serum (Figure 3A-B), which corresponds to the concentration at which complement starts being deposited (Figure S1B). For FcγRIIa-expressing cells, we observed that FcγRIIa gradually increases its interaction with *S. aureus* at low serum concentrations (Figure 3C & S5A). At 0.1% serum however, we observe a decrease in the interaction between bacterial and FcγRIIa-expressing cells. (Figure 3). This effect was due to the presence of the FcγR, as non-expressing control cells displayed no binding across the serum titration (Figure S5A). These data were confirmed in a mAb context, as IgG/IgM depleted serum supplemented with increasing 4497-concentrations enhanced interaction of CR1- and CR3-expressing cells with *S. aureus* (Figure 3D-E). And as in serum, FcγRIIa-expressing cells lost their ability to interact with *S. aureus* at concentrations where complement deposition commences (Figure 3F and S1). This drop in interaction was confirmed with rF1 (Figure S5B).

**Figure 3.**
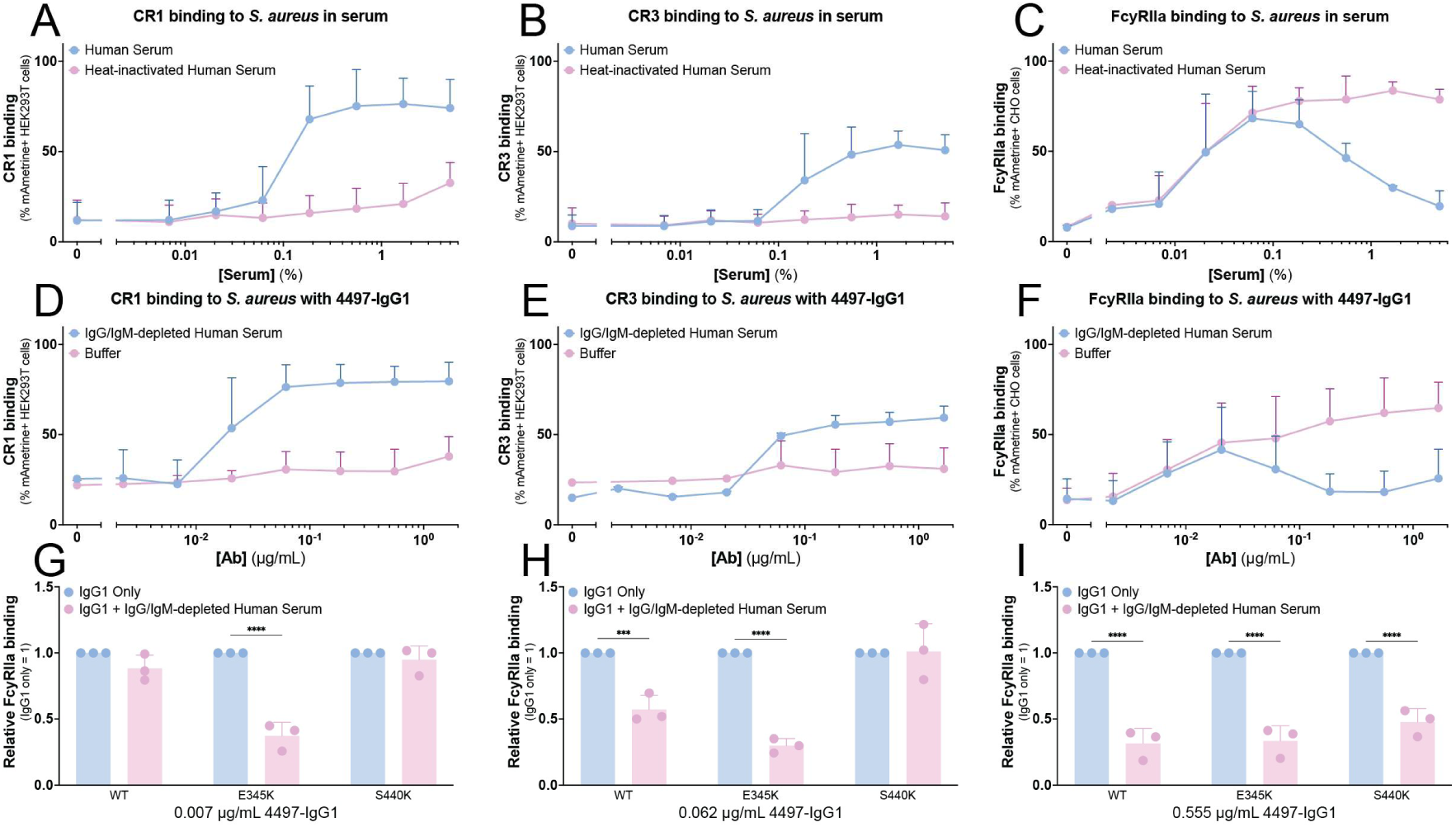
Binding of FcγRIIa to *S. aureus* is reduced by complement deposition. **(A-F)** Binding of *S. aureus* Newman Δ*SpA/Sbi* - mAmetrine to **(A & D)** CR1- or **(B & E)** CR3-expressing HEK293T cells or (**C & F)** FcγRIIa-expressing CHO cells (MOI 10:1), induced by a titration of **(A-C)** (heat-inactivated) human serum, or **(D-F)** mAb 4497 in 1% IgG/IgM-depleted serum or buffer. Binding to cell lines was quantified by flow cytometry and plotted as percentage mAmetrine-positive cells. **(G-I)** Binding of *S. aureus* Newman Δ*SpA/Sbi* - mAmetrine to FcγRIIa-expressing CHO cells (MOI 10:1), induced by mAb 4497 WT, mAb 4497 E345K or mAb 4497 S440K. Bacteria were incubated with **(G)** 0.007 µg/mL, **(H)** 0.062 µg/mL, or **(G)** 0.555 µg/mL mAb in buffer (IgG1 only) or 1% IgG/IgM-depleted serum. Binding to cell lines was quantified by flow cytometry and plotted as percentage mAmetrine-positive cells and plotted as binding normalized to IgG1 only. Data represent mean ± SD of n=3.). Statistical significance was assessed by two-way ANOVA followed by Sidaks’s multiple-comparisons test, ∗∗∗p < 0.001, and ∗∗∗∗p < 0.0001.

Efficient activation of the classical complement pathway requires the organization of antigen-bound IgGs into higher-order Fc assemblies. Following binding to a surface, lateral Fc–Fc interactions promote the formation of noncovalent IgG hexamers, which provide a multivalent docking platform for C1q. Engagement of multiple IgG-Fc domains by the six globular heads of C1q stabilizes binding of the C1 complex and initiates activation of the associated C1r and C1s proteases. IgG hexamerization therefore strongly influences the efficiency of complement deposition while leaving the intrinsic interaction between individual IgG-Fc domains and FcγRs unaffected. Therefore, we asked whether the interference of complement with FcγRIIa binding was related to the capacity of surface-bound IgGs to form hexamers. To test this, we produced two variants of 4497 that alter Fc-dependent hexamerization and C1q recruitment without directly affecting FcγR binding: E345K and S440K.^41–43^ E345K promotes Fc–Fc interactions and IgG hexamerization on the bacterial surface, thereby enhancing C1q binding and complement deposition. In contrast, S440K introduces charge repulsion between adjacent Fc domains, reducing IgG hexamerization and consequently complement activation. Consistent with their expected effects on complement activation, E345K increased whereas S440K decreased the binding of opsonized bacteria to both CR1- and CR3-expressing cells. (Figure S5C-D). For FcγRIIa-expressing cells, we observed that in the presence of complement, the interaction between 4497-E345K-opsonized *S. aureus* is already inhibited at low antibody concentrations compared to 4497-WT (Figure 3G-I and S5E-F), while the opposite phenotype is observed when bacteria were opsonized with 4497-S440K (Figure 3G-I and S5E-F). In all, we show that complement enhances recognition of bacteria via CRs, while simultaneously preventing FcγR binding.

### C1q is responsible for the interference of FcγRIIa:IgG-Fc interaction

Having established that complement prevents access of FcγRIIa to IgG-Fc, we wondered which complement component(s) are responsible for this effect. Since C1q and FcγRs bind to overlapping regions of the Fc tail,^44,45^ we hypothesized that binding of the C1 complex directly interferes with FcγR engagement. Alternatively, covalent deposition of downstream complement components near antigen-bound IgG could sterically restrict FcγR binding. To distinguish between these possibilities, we sequentially added purified complement components according to their order of activation in the complement cascade (Figure 4A). After incubation of *S. aureus* by complement proteins, we let recombinant soluble FcγRIIa-HIS (sFcγRIIa) bind to the opsonized bacteria and measured bound receptor via flow cytometry. When incubating all complement components at 2.5% serum equivalent, just adding C1 and C1 inhibitor was enough to almost fully prevent sFcγRIIa binding (Figure 4B). Titration of C1 in the presence of a constant concentration of 4497 revealed that C4 further reduced sFcγRIIa binding at limiting C1 concentrations (Figure 4C). No sFcγRIIa binding was detected when bacteria were incubated with a non-binding isotype-control antibody (Figure S5G-H). Consistent with the binding experiments, addition of either the complete C1 complex or purified C1q was sufficient to abolish FcγR-mediated phagocytosis of S. aureus (Figure 4D-E). Together, these findings identify C1q as the principal mediator of complement-dependent inhibition of FcγRIIa–IgG-Fc interactions, while C1-dependent C4b deposition can provide an additional inhibitory effect when C1 concentrations are limiting.

**Figure 4.**
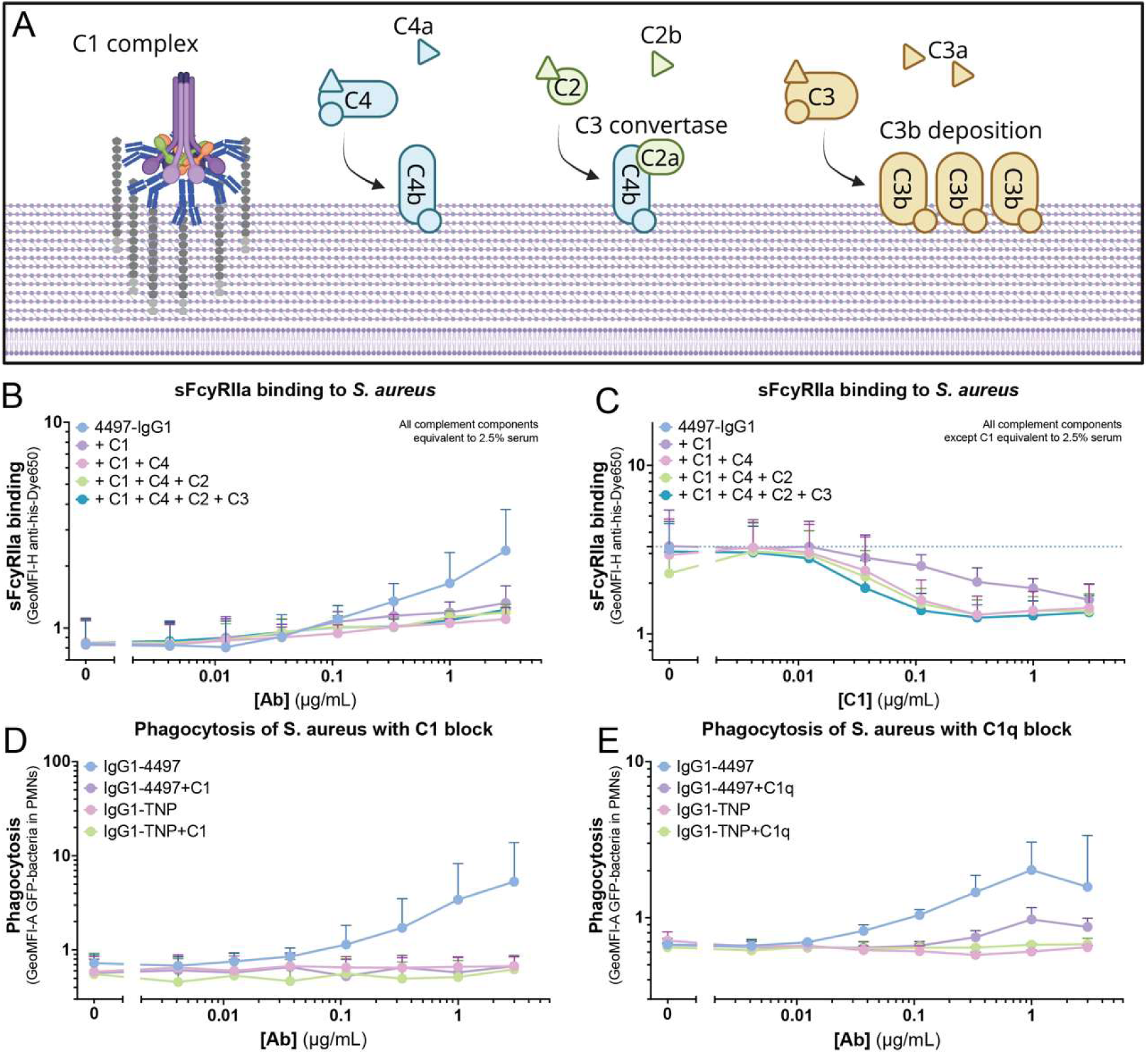
C1q is responsible for the interference of FcγRIIa:IgG-Fc interaction. **(A)** Schematic overview of the first steps of the classical complement pathway, starting with docking of the C1 complex to IgG hexamers, until C3b deposition on the bacterial cell wall. Created in Biorender. **(B)** Binding of soluble FcγRIIa-His to *S. aureus* LAC Δ*SpA/Sbi* - GFP, opsonized with titrated mAb 4497 with complement proteins C1 (+ C1-inhibitor), C4, C2 and C3 added in a stepwise manner at the equivalent of 2.5% serum. **(C)** Binding of soluble FcγRIIa-His to *S. aureus* LAC Δ*SpA/Sbi* - GFP, opsonized with mAb 4497 with complement proteins C1 (+ C1-inhibitor), C4, C2 and C3 added in a stepwise manner. C1 (+ C1-inhibitor) is titrated and the other complement components were added at the equivalent of 2.5% serum. **(B-C)** Binding of sFcγRIIa was quantified by flow cytometry, by measuring GeoMFI of anti-His-Dye650 in the GFP+ bacteria population and data represent mean ± SD of n=3. **(D-E)** Phagocytosis (MOI 10:1, 15 min incubation) of *S. aureus* LAC Δ*SpA/Sbi* - GFP after coincubation with human neutrophils. Bacteria were opsonized with titrated mAb 4497 or isotype control mAb TNP together with **(D)** C1 and C1-inhbiitor or **(E)** C1q, all at 2.5% serum equivalent. **(D-E)** GeoMFI of GFP in the neutrophil population was determined via flow cytometry and data represent mean ± SD of n=3.

### Restoring access to IgG-Fc does not enhance phagocytosis or intracellular killing

Lastly, we aimed to restore access of FcγRs to IgG-Fc to potentially enhance the phagocytic process and boost killing of *S. aureus*. To do this, we made use of a panel of human anti-C1q IgG1s. These antibodies bind the collagen-like region of solid-phase C1q when in complex with antigen-bound IgG, while not binding C1q in solution (Figure 5A).^46^ These antibodies were shown not to influence complement activation, but enhanced FcγR engagement and function through their Fc domains.^46^ However, the authors did not test whether these antibodies are able to boost phagocytosis or intracellular killing in the presence of functioning CRs. First, we screened all 6 anti-C1q mAbs in their ability to rescue FcγR-mediated phagocytosis of *S. aureus* after inhibition with C1. In our model, only 4E6 was able to restore phagocytosis, while the other anti-C1q antibodies did not or only partly restored phagocytosis (Figure 5B). Continuing only with the 4E6 clone, we tested whether this mAb could improve phagocytosis in the presence of all serum components. However, we observed no beneficial effect when adding 4E6; neither at the level of phagocytic uptake (Figure 5C), nor at the level of intracellular killing (Figure 5D). These data show that even when FcγR signaling is restored, these receptors still have a very negligible effect in the presence of complement proteins.

**Figure 5.**
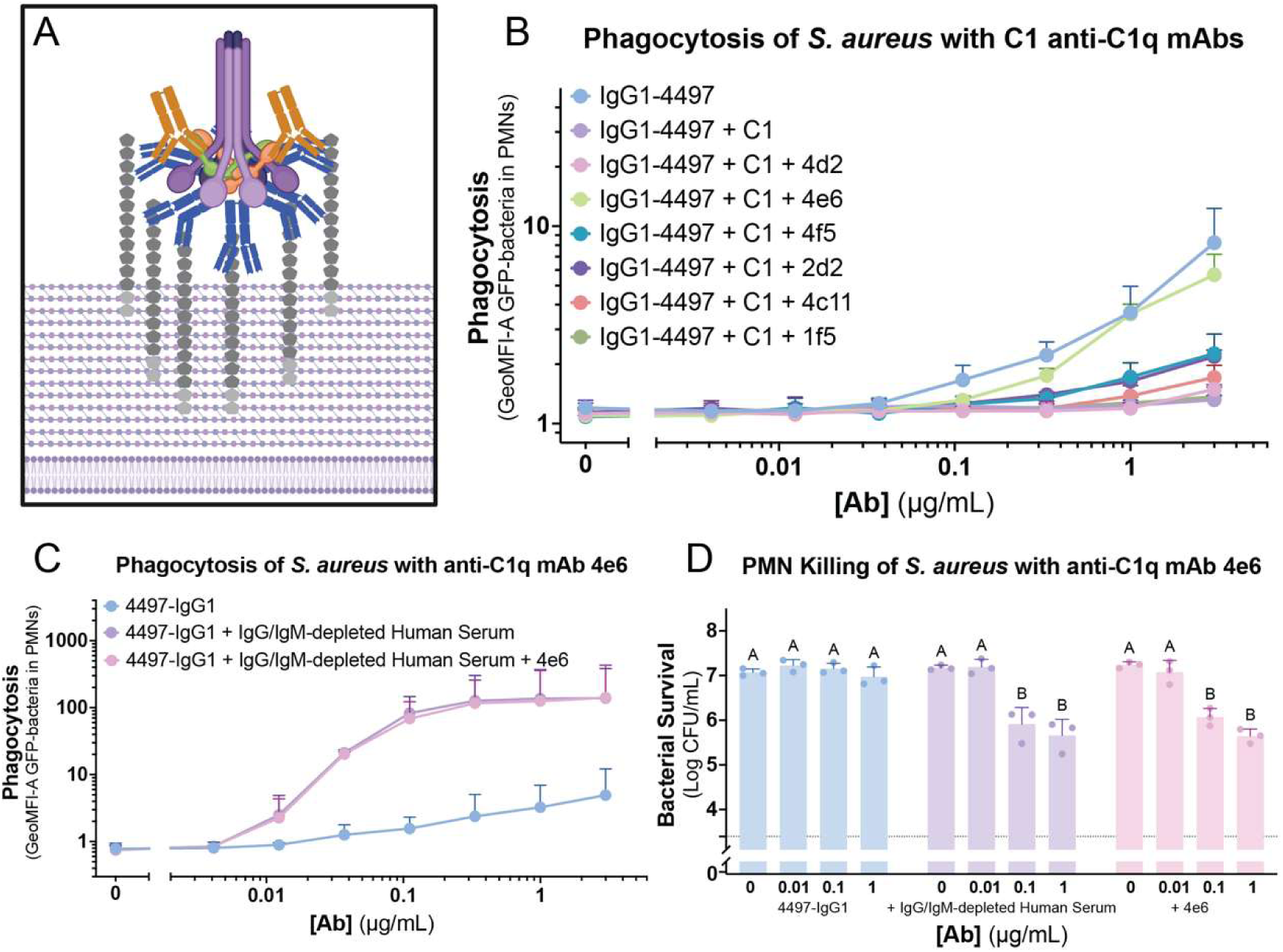
Restoring access to IgG-Fc does not enhance phagocytosis or intracellular killing. **(A)** Schematic overview of anti-WTA mAb (blue) binding WTA on the *S. aureus* cell wall, following by recruitment of the C1 complex (purple). Anti-C1q mAbs (orange) bind C1q only in solid phase and provide exposed Fc-domains. Created in Biorender. **(B)** Phagocytosis (MOI 10:1, 15 min incubation) of *S. aureus* LAC Δ*SpA/Sbi* - GFP after coincubation with human neutrophils. Bacteria were opsonized with titrated mAb 4497 together with C1 and C1-inhibitor at 2.5% serum equivalent and anti-C1q mAbs (3 µg/mL). **(C)** Phagocytosis (MOI 10:1, 15 min incubation) and **(D)** bacterial survival (MOI 1:1, 90 min incubation) of *S. aureus* LAC Δ*SpA/Sbi* - GFP after coincubation with human neutrophils. Bacteria were opsonized with titrated mAb 4497 together with buffer or IgG/IgM-depleted human serum and buffer or anti-C1q mAb 4e6 (3 µg/mL). **(B-C)** GeoMFI of GFP in the neutrophil population was determined via flow cytometry and data represent mean ± SD of n=3. **(D)** Data are visualized as log-transformed CFU/mL values and displayed as mean ± SD of n=3. Statistical significance was assessed by two-way ANOVA followed by Tukey’s multiple-comparisons test. Groups that do not share a letter are significantly different (p < 0.05).

## Discussion

In this study, we show that complement receptors (CRs) are the predominant drivers of antibody-mediated phagocytosis of *S. aureus* and *K. pneumoniae* by human neutrophils when complement is available. Across polyclonal serum, mAbs targeting different bacterial antigens and multiple IgG subclasses, specific inhibition of FcγRs using FLIPr-like generally had little effect on phagocytic uptake or bacterial killing in the presence of complement. Moreover, in the absence of complement, relatively modest FcγR-mediated uptake of *K. pneumoniae* was associated with efficient bacterial killing. Mechanistically, we demonstrate that complement deposition, and particularly C1q binding to IgG-Fc, restricts FcγRIIa access to surface-bound IgG. Thus, complement not only generates additional ligands for CRs but also shifts antibody-mediated recognition away from FcγRs.

FcγRs and CRs have historically been associated with distinct modes of phagocytosis and downstream signaling, although more recent work indicates substantial morphological and functional overlap between these pathways.^47–50^ Functional crosstalk has also been reported: CR3 and FcγRIII can cooperate in the induction of the neutrophil oxidative burst, while FcγRIIa and FcγRI can differentially regulate CR3-mediated phagocytosis in macrophages.^51,52^ Our findings add another layer to this interplay. When both antibody and complement opsonins are present on the same bacterial surface, the pathways may not function independently or additively because recruitment of complement can physically reduce the availability of IgG-Fc for FcγR engagement.

The relative contributions of these pathways are likely to depend on the antigen and its organization on the bacterial surface. Most antibodies tested in this study recognize abundant and repetitive glycan structures, including wall teichoic acid on *S. aureus* and the O-antigen of *K. pneumoniae*. Such high-density antigens are favourable for lateral Fc–Fc interactions, IgG hexamerization and C1q recruitment. By contrast, the MrkA-targeting antibody showed a less pronounced dominance of CR-mediated phagocytosis, suggesting that differences in antigen density, distribution or accessibility may influence the extent to which complement interferes with FcγR binding.

These findings have important implications for the development of therapeutic antibodies against bacterial pathogens. Antibody engineering frequently aims to enhance either FcγR engagement or complement activation through modifications of the Fc domain.^34,42,53–55^ Our results indicate that simultaneously optimizing both functions may not necessarily produce additive neutrophil responses. For antibodies intended to promote neutrophil-mediated clearance in complement-rich environments, efficient complement activation and CR engagement may therefore be more important than increasing intrinsic FcγR affinity. In Gram-negative bacteria susceptible to terminal complement-mediated lysis, enhanced complement activation could additionally promote direct killing through the membrane attack complex, although the *K. pneumoniae* strains used here were resistant to membrane attack complex-mediated killing and our experiments therefore do not directly assess this potential benefit. Conversely, preserving FcγR accessibility may be advantageous when FcγR-mediated uptake or signalling produces a more favourable outcome than CR-mediated uptake. The anti-C1q antibody 4E6 restored FcγR-mediated phagocytosis after C1-dependent inhibition, demonstrating that complement-induced Fc masking can, in principle, be reversed. However, restoring FcγR engagement did not further enhance uptake or killing of *S. aureus* in the presence of intact complement, indicating that increased FcγR availability was not beneficial in this experimental setting. Nevertheless, this strategy could have value for pathogens for which CR-mediated uptake is inefficient or promotes intracellular persistence. For example, CR3-dependent uptake of Opa-negative *N. gonorrhoeae* by neutrophils has been associated with limited oxidative-burst induction, delayed phagosome maturation and bacterial survival.^56^ Although it remains unknown whether redirecting gonococcal uptake towards FcγRs would improve killing, this example illustrates how the receptor pathway used for internalization can influence pathogen fate.

The competition between complement and FcγRs may also be relevant beyond bacterial immunity. Many antitumor antibodies are designed to activate complement while simultaneously recruiting FcγR-expressing natural killer cells or macrophages. Complement deposition on rituximab-coated tumour cells has previously been shown to reduce the interaction between antibody Fc domains and FcγRIIIa on natural killer cells, thereby limiting antibody-dependent cellular cytotoxicity.^57^ This closely resembles the Fc-masking mechanism observed here and indicates that complement-enhancing modifications may reduce FcγR-dependent functions when both pathways act on the same target. In viral infections, inhibition of FcγR engagement can instead be protective. Antiviral antibodies can facilitate infection of FcγR-expressing cells through antibody-dependent enhancement. C1q has been shown to restrict antibody-dependent enhancement of flavivirus infection in an IgG subclass-dependent manner.^58^ Although viral entry and bacterial phagocytosis have different consequences, these observations support a broader model in which C1q binding can regulate the accessibility and function of antibody Fc domains. For bacteria, this redirects recognition towards CRs; for tumours, it may reduce FcγR-mediated cytotoxicity; and for viruses, it may prevent harmful FcγR-dependent entry.

The physiological relevance of FcγR-mediated phagocytosis will also depend on local complement availability. Complement proteins are abundant in the circulation, but their concentrations and activity can be lower at mucosal surfaces and in certain tissue compartments.^59,60^ FcγRs may therefore become more important in complement-limited niches or during infection with pathogens that efficiently evade complement deposition. Nevertheless, we observed impaired FcγRIIa binding at serum concentrations representing only a small fraction of normal circulating levels, suggesting that complete complement depletion is not required for this effect. In addition to complement availability, antigen density, antibody subclass, Fc organization, bacterial immune-evasion mechanisms and inflammation-dependent changes in receptor expression are all likely to shift the balance between FcγR- and CR-mediated recognition.

Several limitations should be considered. Our functional experiments were performed with neutrophils from healthy donors under controlled in vitro conditions and included only two bacterial species. Most mAbs recognized abundant glycan epitopes, limiting the extent to which the results can be generalized to sparsely distributed protein antigens. Furthermore, FcγRIIa accessibility was examined using soluble receptor and receptor-overexpressing cell lines, which isolate individual receptor interactions but do not fully reproduce the receptor density, mobility and signalling environment of primary phagocytes. Finally, although our data identify C1q as the principal inhibitor of FcγRIIa binding, the precise spatial organization of C1q, C4b and C3 fragments around surface-bound IgG remains unresolved.

In conclusion, our findings identify CRs as the principal drivers of antibody-mediated phagocytosis by human neutrophils in the presence of complement and reveal that C1q directly restricts access of FcγRIIa to IgG-Fc. Rather than functioning as fully independent and additive pathways, FcγRs and complement can compete for the functional use of surface-bound antibodies. This framework provides a mechanistic basis for understanding how antibody specificity, Fc properties, complement availability and effector-cell identity together determine antibody function. These factors should be considered when designing therapeutic antibodies intended to recruit complement, FcγRs or both.

## Materials and Methods

### Human material

In accordance with the declaration of Helsinki, human blood was isolated after informed consent from each donor. Approval was obtained from the Medical Ethics Committee of the University Medical Center Utrecht (METC protocol 07-125/C, approved March 1, 2010).

### Isolation of human neutrophils

Neutrophils were isolated from healthy donor blood, as described before.^61^ In short, whole blood was collected in sodium heparin 9 mL tubes (Greiner). Density gradients were prepared by pipetting 10 mL Ficoll (GE Healthcare, 17544203) on top of 12 mL Histopack (density 1.119; Sigma, 11191-100mL). After layering 25 mL blood diluted 1:1 (v/v) with PBS on top of the Histopack/Ficoll stack, the gradients were centrifuged for 20 min at 20°C at 390 x g without brake. Afterwards, the upper plasma and PBMC ring were discarded and the layer containing neutrophils was carefully collected and washed with RPMI (ThermoFisher/Gibco) supplemented with 0.05% human serum albumin (HSA) by centrifugation for 10 min at 4°C at 249 x g. Erythrocytes were lysed by shocking the pallet hypotonically with 9 mL cold deionized H_2_O for 30 sec, after which osmolarity was restored by adding 1 mL 10x PBS. After another washing step for 10 min at 4°C at 249 x g, cells were resuspended in RPMI-HSA at 1 x 10^7^ neutrophil / mL.

### Bacterial strains and culture conditions

Table 1 depicts a list of all used bacterial strains and their growth conditions. GFP and mAmetrine were expressed as described before.^61^ For all experiments, a bacterial colony was inoculated from a plate into medium and left overnight shaking at 37°C. The next day, bacteria were sub-cultured in fresh medium and grown shaking 37°C until mid-log (OD_600_ 0.4-0.6), after which the bacteria were washed and resuspended to OD_600_ = 1 in RPMI-HSA and diluted as required for subsequent experiments.

**Table 1.** Bacterial species and strains used.

| Bacterial strain | Plate type | Growth medium | Antibiotic | Source |
| --- | --- | --- | --- | --- |
| *S. aureus* LAC<br>$\Delta spa/sbi$ GFP | Todd Hewitt<br>Agar | Todd Hewitt<br>Broth | 10 µg/mL<br>Chloramphenicol | This study |
| *S. aureus* LAC<br>$\Delta spa/sbi$ | Todd Hewitt<br>Agar | Todd Hewitt<br>Broth | | <sup>67</sup> |
| *S. aureus*<br>Newman<br>$\Delta spa/sbi$<br>mAmetrine | Todd Hewitt<br>Agar | Todd Hewitt<br>Broth | 10 µg/mL<br>Chloramphenicol | This study |
| *K. pneumoniae*<br>*KpnO1* GFP | LB Agar | Lysogeny Broth | 10 µg/mL<br>Gentamycin | This study |
| *K. pneumoniae*<br>*KpnO1* | LB Agar | Lysogeny Broth |  | <sup>24</sup> |
| *K. pneumoniae*<br>*Kpn570.879* GFP | LB Agar | Lysogeny Broth | 10 µg/mL<br>Kanamycin | This study |
| *K. pneumoniae*<br>*Kpn570.879* | LB Agar | Lysogeny Broth |  | <sup>68</sup> |
| *K. pneumoniae*<br>*Kpn209R* | LB Agar | Lysogeny Broth |  | <sup>69</sup> |

### Production of human monoclonal antibodies

IgG1-4 were produced as described in detail before.^62^ In short, VH and VL sequences (Table 2) were cloned in pcDNA34 vectors (ThermoFisher Scientific) and transfected into Expi293F cells (ThermoFisher Scientific). The E345K and S440K mutations were introduced in the heavy chain expression vectors using gene synthesis (Integrated DNA Technologies). 5 days after transfection, supernatant was collected and dialyzed against PBS. Next the mAbs were purified using an Äkta Pure Sytem (GE Healthcare). For IgG mAbs, a hiTrap Protein G high performance column (GE healthcare) was used. Lastly, IgG mAbs were isolated on a Superdex 200 Increase 10/300 GL to ≥95% purity. Antibodies were stored at 4°C.

**Table 2.**
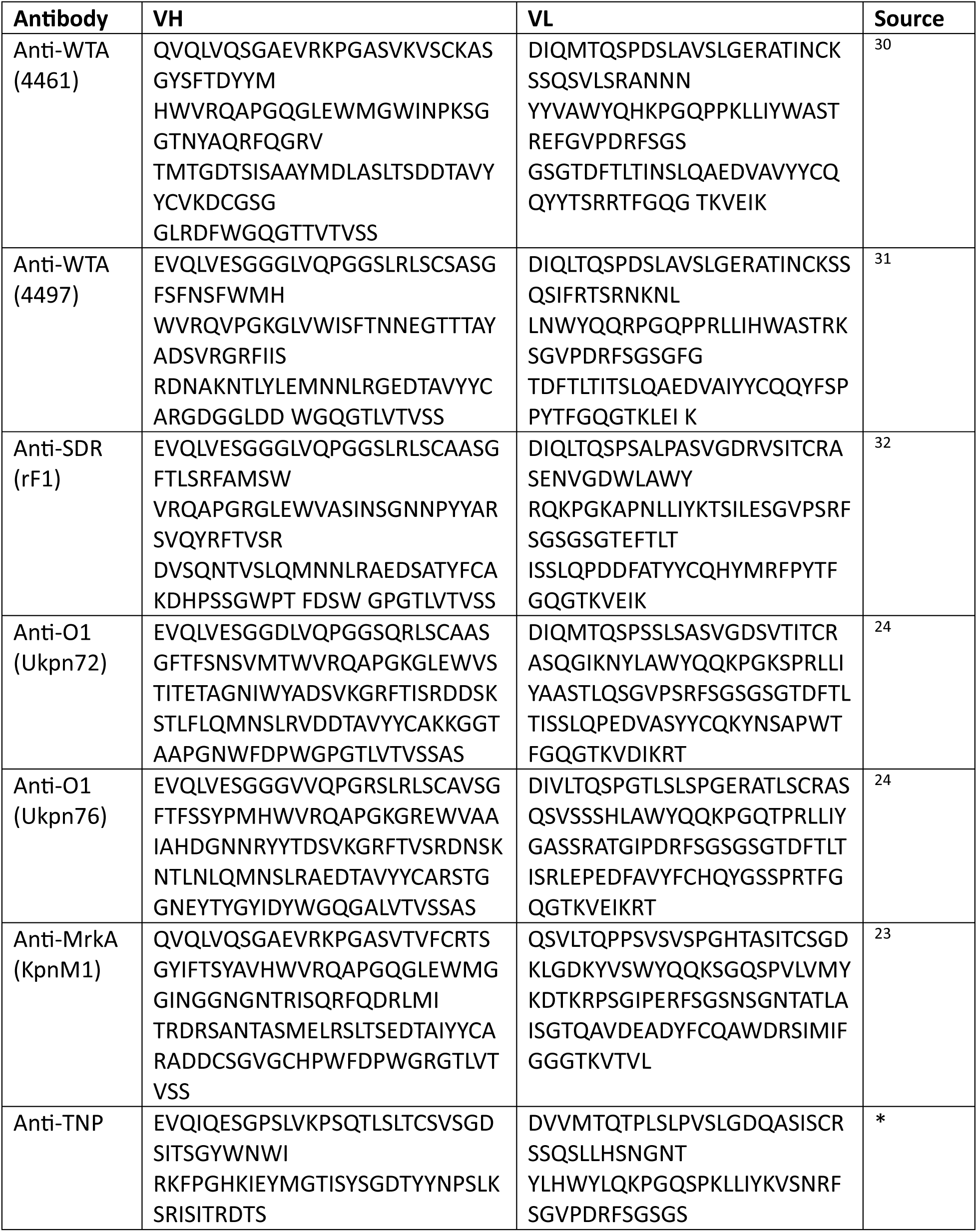
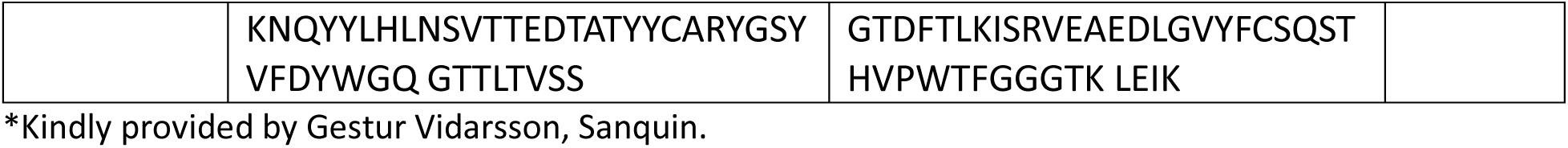
Variable and constant heavy chain and light chain amino acid sequences used for antibody production.

### Preparation of normal human serum and IgG/IgM depletion

Fresh blood was collected in tubes not containing anticoagulant (Greiner), allowing to clot for 30 min at room temperature. Centrifugation for 10 min at 4°C at 2075 x g resulted in serum as supernatant. To correct for donor variability, sera from 20 healthy donors was pooled and aliquoted at -80°C. Complement components were thermally inactivated with a 30 min incubation at 56°C.

Serum was depleted of IgG and IgM as described before.^63^ In short, 1M NaCl was added to the serum beore using an Äkta Pure Sytem to capture IgG, using hiTrap Protein G High Performance column (GE Healthcare). Similarly, IgM was depleted using Capture Select IgM Affinity Matrix (Thermo Fisher Scientific). All batches depleted serum were tested for complement levels and activity in classical and alternative pathway hemolytic assays.

### Antibody binding assay

Bacteria were cultured as described above and incubated for 30 min shaking at 4°C at a bacterial concentration of OD_600_ = 0.025 co-incubated with specific antibody concentrations indicated per experiment. After washing with RPMI-HSA via a 10 min centrifugation at 2.400 x g, bacteria were resuspended in 3 µg/mL Goat-anti-human-kappa-AF647 (Southern Biotech, 2060-31) or 3 µg/mL Goat-anti-human-IgG-AF647 (Southern Biotech, 2040-31) detection antibody for 30 min shaking at 4°C. After incubation, bacteria were washed again and resuspended in RPMI-HSA + 1% paraformaldehyde (PFA). Bacteria were then analyzed by flow cytometry on a MACSQuant X (Miltenyi Biotec). After gating GFP+ events based on SSC-H and FITC-H to distinguish bacteria from background, median Fluorescent Intensity (MFI) of the APC-H channel was used as a parameter for antibody binding.

### Complement deposition assay

Bacteria were cultured as described earlier and incubated for 15 min shaking at 4°C at a bacterial concentration of OD_600_ = 0.05 or OD_600_ = 0.025 when co-incubated with an antibody or (heat-inactivated) serum titration respectively, specific concentrations indicated per experiment. When mAbs were used in an experiment, complement was supplied by incubating for 15 min shaking at 37°C in IgG/IgM-depleted serum equivalent to 1% serum for S. aureus and 5% for K. pneumoniae, at which point the bacterial concentration was diluted to OD_600_ = 0.025. After washing with RPMI-HSA via a 10 min centrifugation at 2.400 x g, bacteria were resuspended in 3 µg/mL polyclonal rabbit-*anti*-human-C1q-FITC (Dako, F0254) or 3 µg/mL Goat-*anti*-human-C3b-AF647 (Clone bH6^64^, randomly labeled with NHS-Alexa Fluor 647 (Invitrogen, A20006)) detection antibody for 30 min shaking at 4°C. After incubation, bacteria were washed again and resuspended in RPMI-HSA + 1% paraformaldehyde (PFA). Bacteria were then analyzed by flow cytometry on a MACSQuant X. After gating GFP+ events based on an SSC-H vs FITC-H to distinguish bacteria from background, MFI of the APC-H channel was used as a parameter for C3b deposition. In experiments with serum, non-fluorescent bacteria were measured, and the FITC-H channel and APC-H channels were used to simultaneously measure C1q and C3b deposition respectively.

### Phagocytosis flow cytometry assay

Phagocytosis was measured by flow cytometry as described previously.^61^ Briefly, GFP-expressing bacteria were cultured and incubated with titrations of (heat-inactivated) serum, or antibody titrations followed by 1% (*S. aureus*) or 5% (*K. pneumoniae*) IgG/IgM-depleted serum incubation, as described for the complement deposition assay. Subsequently, freshly isolated neutrophils were added and incubated shaking for 15 min at 37°C at a multiplicity of infection (MOI) of approximately 10 bacteria per neutrophil. To block FcyRs on neutrophils, neutrophils were pre-incubated on ice for 15 min with FLIPr-like (3 µg/mL), which was expressed and purified as described before.^29^ Samples were then fixed in cold RPMI-HSA containing 1% PFA and analyzed by flow cytometry using a MACSQuant X. neutrophils were identified based on forward scatter (FSC-A) and side scatter (SSC-A) characteristics. Neutrophils were gated using FSC-A versus FSC-H. Phagocytosis was assessed by determining GFP fluorescence in the FITC-A channel, and neutrophils were classified as GFP-positive or GFP-negative based on this signal. Phagocytic uptake was quantified as both the median fluorescence intensity (MFI) of the total neutrophil population and the percentage of GFP-positive neutrophils.

### Widefield microscopy analysis of phagocytosis

Phagocytosis was induced as described above. After fixation, 3 µg/mL WGA AF647 (ThermoFisher Scientific, W32466) was used to stain membranes of neutrophils as well as cell walls of non-phagocytosed *S. aureus* bacteria for 5 min at 4°C shaking. Next, samples were washed by centrifuging at 300 x g for 5 min and resuspending in 10 µL RPMI-HSA. 10 µL sample was then immobilized by loading them on solidified agar pads consisting of 1% low-melting (SeaPlaque) agarose (Lonza) in PBS and dried for 10 min before covering the sample with a 22 x 22 mm coverslip (Superior Marienfeld). A Leica SP5 microscope equipped with a HC PL APO100x/1.40 OIL PH3 objective. WGA-AF647 and GFP-labeled bacteria were detected using the Y5 (excitation 620/60 nm, emission 700/75 nm) and GFP (excitation 470//40 nm, emission 525/50 nm) filter cubes respectively. Next to the fluorescent channels, phase contrast images were included in all experiments. Manually in ImageJ, the number of intracellular and extracellular *S. aureus* bacteria were counted for each neutrophil that was completely in frame. Bacteria were classified as phagocytosed when they were WGA negative and localized inside the neutrophil. Bacteria were classified as non-phagocytosed when they were WGA positive and were attached to a neutrophil. Using these counts, net phagocytic uptake (number of *S. aureus* inside neutrophil) and phagocytosis efficiency (Percentage of *S. aureus* inside neutrophil).

### Bacterial colony forming unit assay

Opsonophagocytic killing of was measured by culturing bacteria as described above, but washing and resuspending in HBSS (Gibco) with 0.1% HSA instead of RPMI-HSA. Bacteria were incubated at OD_600_ = 0.03 with per experiment indicated titrations of (heat-inactivated) serum, or antibody titrations followed by 1% IgG/IgM-depleted serum at 30 min at 37°C. Then, neutrophils were added at an MOI of 1, resulting in bacterial OD_600_ = 0.0085. Bacteria and neutrophil co-incubated shaking for 90 min at 37°C with 5% CO_2_. After 90 min, neutrophil were lysed by adding 0.3% saponin (Sigma) in MiliQ water for 5 min on ice. Bacteria were then serial diluted in steps of 1:10 in PBS and plated on appropriate agar plates without antibiotics. After overnight incubation at 37°C, colonies were counted as a parameter of neutrophil killing.

### Culturing of hFcγR CHO and hCR1 and hCR3 HEK cell lines

CHO cells expressing human FcyRIIA were generated at the University of Erlangen-Nürnber laboratory (Lux et al. 2013). These cells were grown in RPMI medium supplemented with 10% FCS, 2 mM glutamine, 1 mM sodium pyruvate, 0.1 mg/mL pen/strep, 0.1 mM non-essential amino acids and 0.05 mg/mL G418 (Gibco) at 37°C with 5% CO_2_. Cells were detached by brief trypsinization, washed and resuspended in RPMI-HSA at 7.5x10^6^ cells/mL.

To stably express CR3 in HEK293T cells, we cloned CD18 (NM_00211.4) and CD11b (NM_001145808.1) cDNA in bicistronic expression vectors pIRESpuro3 and pIREShyg3 vector (Clontech) respectively. First, the pIRESpuro3/CD18 construct was transfected using Lipofectamin 2000 (Life Technologies) after which cells were put on selection pressure using 1 µg/mL puromycin in DMEM (Gibco, 11995) with 10% FBS. A few weeks later, puromycin resistant cells were transfected with the pIREShyg3/CD11b construct and put under selection pressure with 250 µg/mL hygromycin. Stable CR3 expression was verified with flow cytometry analysis using *anti-* CD11b-APC (BD, clone ICRF44) and *anti*-CD18-FITC (BD, clone 6.7).

For stable expression of human CR1 in HEK293T cells, we used a lentiviral expression system. Therefore, a codon-optimized CR1 gene (NM_000651) was cloned in the dual promoter lentiviral vector, RP137 (BIC-PGK-Zeo). This vector was constructed by replacing the Zeo-T2a-mAmetrine cassette from the BIC-PGK-Zeo-T2a-mAmetrine (RP172) vector^65^ with a Zeocin resistance gene. Virus was produced in 24-well plates using standard lentiviral production protocols and the third-generation packaging vectors pMD2G-VSVg, pRSV-REV, and pMDL/RRE. Briefly, 0.25 µg lentiviral vector and 0.25 µg packaging vectors were cotransfected in HEK293T cells by using 1.5 ul Mirus LT1 tranfection reagent (Sopachem, Ochten, The Netherlands). After 72 hours, 100 µl unconcentrated viral supernatant adjusted to 8 mg ml-1 polybrene was used to infect ∼50,000 HEK293T cells by spin infection at 1,000g for 2 h at 33°C. Three days after infection, cells were put under selection pressure using 500 µg/ml zeocin. Stable CR1 expression was verified with flow cytometry using mouse anti-human CR1(CD35)-PE (BD, clone E11) mAmetrine-expressing *S. aureus* bacteria were incubated with titrations of (heat-inactivated) serum or antibody titrations followed by 1% IgG/IgM-depleted serum for 15 min shaking at 37°C to induce antibody and/or complement opsonization. Then, opsonized bacteria were co-incubated with FcγRIIa, CR1 or CR3 expressing cells for 30 min shaking at 37°C, washed once and then measured via flow cytometry (BD FACSverse). Cells were gated via forward scatter (FSC-A) and side scatter (SSC-A), after which the percentage of mAmetrine+ cells (FITC-A) was used as a parameter of receptor binding.

### Purified complement component assay and sFcγRIIa binding

The purified classical pathway assay was performed as described before ^62^, only using complement components C1-complex (Complement Technology, A098), C1 inhibitor (Complement Technology, A140), C4 (Complement Technology, A105), C2 (Complement Technology, A112) and C3 (Isolated from human plasma). Purified complement components were used in concentrations in physiological ratio as present in normal healthy human serum. The following concentrations were regarded as 100% serum: C1-complex; 135 µg/mL, C1-inhibitor; 20 µg/mL, C4; 70 µg/mL, C2; 1250 µg/mL, and C3; 400 µg/mL. Bacteria at OD_600_ = 0.04 were incubated as described above and incubated with an antibody titration and C1-complex while shaking for 15 min at 4°C. Next, all subsequent complement components were added as indicated to achieve a final complement protein concentration of 2.5% serum equivalent and a bacterial OD_600_ = 0.01, incubated shaking for 15 min at 37°C. Bacteria were washed once before continuing with subsequent experiments. To measure sFcγRIIa binding to bacteria opsonized by complement components added in a step-wise fashion, bacteria were incubated with 5 µg/mL sFcγRIIa a-HIS (produced in Expi cells) and incubated shaking for 30 min at 4°C. After one washing step, bacteria were resuspended in 3 µg/mL goat-*anti*-HIS-dy650 detection antibody (LSBIO, LS-C181054) and incubated shaking for 30 min at 4°C. Lastly, bacteria were fixed using RPMI-HSA containing 1% PFA and analyzed by flow cytometry using a MACSQuant X. sFcγRIIa a binding was quantified by measuring MFI of Dy650 (APC-H) in GFP+ events (FITC-H).

To measure the effect of C1 on FcyR-mediated phagocytosis, bacteria were cultured as above and incubated with an antibody titration together with C1-complex (2.5% serum equivalent) or an RPMI control incubated shaking 15 min at 37°C, after which C1-inhibitor (2.5% serum equivalent) was added and incubated shaking 15 min at 37°C. Next, neutrophils were added as in previously described phagocytosis assays and samples were fixed and analyzed as described above.

### Anti-C1q antibody experiments

The anti-solid-phase C1q antibodies (REF) were provided by Prof. Trouw, Leiden University. Phagocytosis and colony forming unit assays involving anti-C1q antibodies were performed as described earlier. Anti-C1q antibodies were added at a concentration 3 µg/mL simultaneously with the serum or purified complement components.

### Sytox inner membrane permeabilization assay

Sytox assays were performed like decribed before.^66^ Briefly, bacteria were grown to midlog phase in LB medium and resuspended to OD_600_ = 1 in RPMI-HSA buffer. Bacteria were then diluted to OD_600_ = 0.1 with a 10, 5, 2.5, 1.25, 0.63 or 0% pooled normal human serum, together with 1 µM Sytox Green Nucleic Acid stain (Thermofisher). Fluorescence was measured in a microplate reader (CLARIOstar, Labtech) at 37°C under non-shaking conditions for 90 minutes.

### Statistical analysis

Flow cytometry data were analyzed in FlowJo V10 (FlowJO LLC). Microscopy data were analyzed in ImageJ v1.47c. All graphs were made in Graphpad Prism 10 software, which was also used for all statistical analysis. For each figure the used statistical test is indicated. In all statistical tests, p-values of <0.05 were considered statistically significant.

## Acknowledgments

The authors greatly thank prof. Falk Nimmerjahn (University of Erlangen-Nürnberg) for providing FcγRIIa-expressing CHO cell lines. This project received funding from the Netherlands Organisation for Scientific Research (NWO) through a Vici Grant (no. 09150182210011 to S.H.M.R), the European Research Council (ERC) under the European Union’s Horizon 2020 research and innovation program (grant agreement no. 101001937, ERC-ACCENT, to S.H.M.R.). Also, this project was financially supported by grants from Health∼Holland (LSHM17026) and the Research along Routes by Consortia program [NWA-ORC] (DARTBAC, NWA 1292.19.354).

## Author Contributions

Max L.B. Grönloh: Conceptualization, Formal Analysis, Investigation, Methodology, Supervision, Visualization, Writing - Original draft, Writing - review & editing

Lisanne de Vor: Conceptualization, Formal Analysis, Investigation, Methodology, Visualization, Writing - original draft, Writing - review & editing

Seline Zwarthoff: Conceptualization, Investigation, Methodology, Writing - review & editing

Jana Venema: Formal Analysis, Investigation, Validation, Visualization, Writing - review & editing

Rosemieke Smits: Formal Analysis, Investigation, Validation, Visualization, Writing - review & editing

Stan Baijens: Investigation, Methodology, Writing - review & editing

Maartje Ruyken: Investigation, Validation, Writing - review & editing

Piet Aerts: Investigation, Methodology, Writing - review & editing

Carla J.C. de Haas: Investigation, Methodology, Writing - review & editing

Kok van Kessel: Investigation, Methodology, Writing - review & editing

Leendert A. Trouw: Resources, Writing - review & editing

Bart Bardoel: Conceptualization, Supervision, Writing - review & editing

Suzan Rooijakkers: Conceptualization, Funding acquisition, Supervision, Writing - review & editing

